# Motor Imagery Affects both Cortical and Spinal Circuitry: A Transcranial and Transspinal Magnetic Stimulation Study

**DOI:** 10.1101/2025.11.06.686943

**Authors:** Asma Benachour, Nikolay Syrov, Mikhail Lebedev

## Abstract

Motor imagery (MI), the mental rehearsal of movement without physical execution, is a key technique in brain-computer interfaces (BCIs) for voluntarily eliciting cortical modulations. Beyond cortical effects, MI could also modulate spinal cord processing, which offers additional potential for neurorehabilitation in conditions like spinal cord injury (SCI) and stroke, where BCIs are used for therapy. To investigate the interactions of MI with both the cortex and the spinal cord, we employed both transcranial magnetic stimulation (TMS) and trans-spinal magnetic stimulation (TSMS). With proper coil orientation, TSMS elicited short- and long-latency motor evoked potentials (MEPs) in forearm muscles and lateralized evoked potentials in the cortex. MI modulated both TMS-induced and TSMS-induced cortical responses and MEPs. This demonstration of MI affecting both cortical and spinal circuitry underscores its potential as a powerful strategy for BCI-driven neurorehabilitation, including pairing MI with magnetic stimulation.

## Introduction

Motor Imagery (MI), the mental rehearsal of movement without execution [1], is a well-established paradigm for promoting neuroplasticity in motor rehabilitation [2] and for being used as a control signal in Brain-Computer Interfaces (BCIs) [3]. MI engages a cortical network largely overlapping with that of physical movement, including the primary motor cortex [4], premotor areas [5], the supplementary motor area [6], and posterior parietal cortices [7].

Transcranial magnetic stimulation (TMS) has been proven effective in probing the central nervous system during MI. Studies combining TMS with EEG showed an increase in cortical activity during MI compared to rest, though not to the same extent as during physical movements. Moreover, it has been demonstrated that MI enhances MEPs in muscles associated with the imagined movements, suggesting modulation of cortico-spinal excitability [8–11]. In addition to the effects revealed using TMS, MI influences spinal reflexes, including enhancement of the H-reflexes in limb muscles [12–13], increased spinal reflexes during transcutaneous spinal cord stimulation [14], and larger F-wave amplitudes representing antidromic responses reflecting motoneuron excitability [15].

Thе effects of MI on the spinal circuitry are particularly relevant in the context of non-invasive spinal cord stimulation techniques, such as transcutaneous electrical and magnetic stimulation, which are becoming popular as rehabilitation tools for spinal cord injury (SCI) [16–17].

Spinal excitability can also be tested with transspinal spinal magnetic stimulation (TSMS), a non-invasive technique that involves the delivery of brief magnetic pulses to the spinal cord. While research combining MI with TSMS remains limited; in the BCI domain, a recent study demonstrated a non-invasive brain-spine interface (BSI) where EEG signals from MI of leg movements continuously controlled TSMS over the lumbar spine, eliciting muscle activation in the tibialis anterior muscle [18]. Although these approaches highlight the feasibility of integrating MI with spinal stimulation, applications in clinical populations are still scarce. It has been suggested that combining MI with TSMS could enhance plasticity and improve rehabilitation outcomes in SCI [19–21]. In the current study, we employed TSMS and TMS to probe the effects of MI at the spinal and cortical levels. We adjusted coil parameters for this paradigm and found that the figure-eight coil achieved good focality and laterality when positioned at a 45-degree angle over the C6–C7 vertebrae. With this approach, we demonstrated the modulatory effects of MI on the responses elicited by the stimulation of both spinal cord and cortex.

## Material and methods

Ethical approval was obtained from the Ethical Committee of the Skolkovo Institute of Science and Technology (No. 11 dated October 17, 2023), and the study was conducted following the principles of the Declaration of Helsinki. All participants provided written informed consent prior to their participation in this study.

### Experiment 1 (Pilot Study)

The pilot experiment aimed to investigate the feasibility of using the TMS figure-eight coil to stimulate the spinal cord and elicit lateralized MEPs in the ipsilateral arm and hand muscles. A total of 10 healthy participants (10 males, mean age 26.6;19 to 34 years old, one left-handed) were recruited, all of whom met inclusion criteria of the absence of neurological, motor, psychiatric and cognitive impairments. The exclusion and inclusion criteria conformed to the general TMS criteria [19]. Data from 2 participants was excluded due to issues related to data file acquisition.

### TMS Experimental Design and Setup

#### Spinal Cord Stimulation

The experimental design comprised four distinct stimulation conditions targeting two regions of interest: the right and left sides of the spinal cord at the C6-C7 vertebral level (corresponding to the C5 and C6 nerve roots). For each stimulation site, two experimental conditions were tested: resting and MI of closing and opening of the right hand.

Left-side stimulation was used to investigate the side-specificity of TSMS effects with and without MI of the right-hand.

TMS pulses were delivered at the C6-C7 level using a figure-eight coil (70 mm diameter; Nexstim, Finland) positioned at a 45-degree angle with respect to the spinal cord (see Fig. 1 for an example of right-hand MEP induction). The C6-C7 vertebral level was identified using established anatomical landmarks, the hotspot was found by moving slightly the coil around the anatomical area and tracking the MEPs responses. The resting motor threshold (RMT), defined as the stimulation amplitude inducing MEPs >50 µV in 5 out of 10 consecutive pulses, was determined separately for each spinal cord site immediately prior to the corresponding stimulation condition. Stimulation intensity was set to 110% of RMT. The order of conditions was randomized across participants.

**Figure 1.**
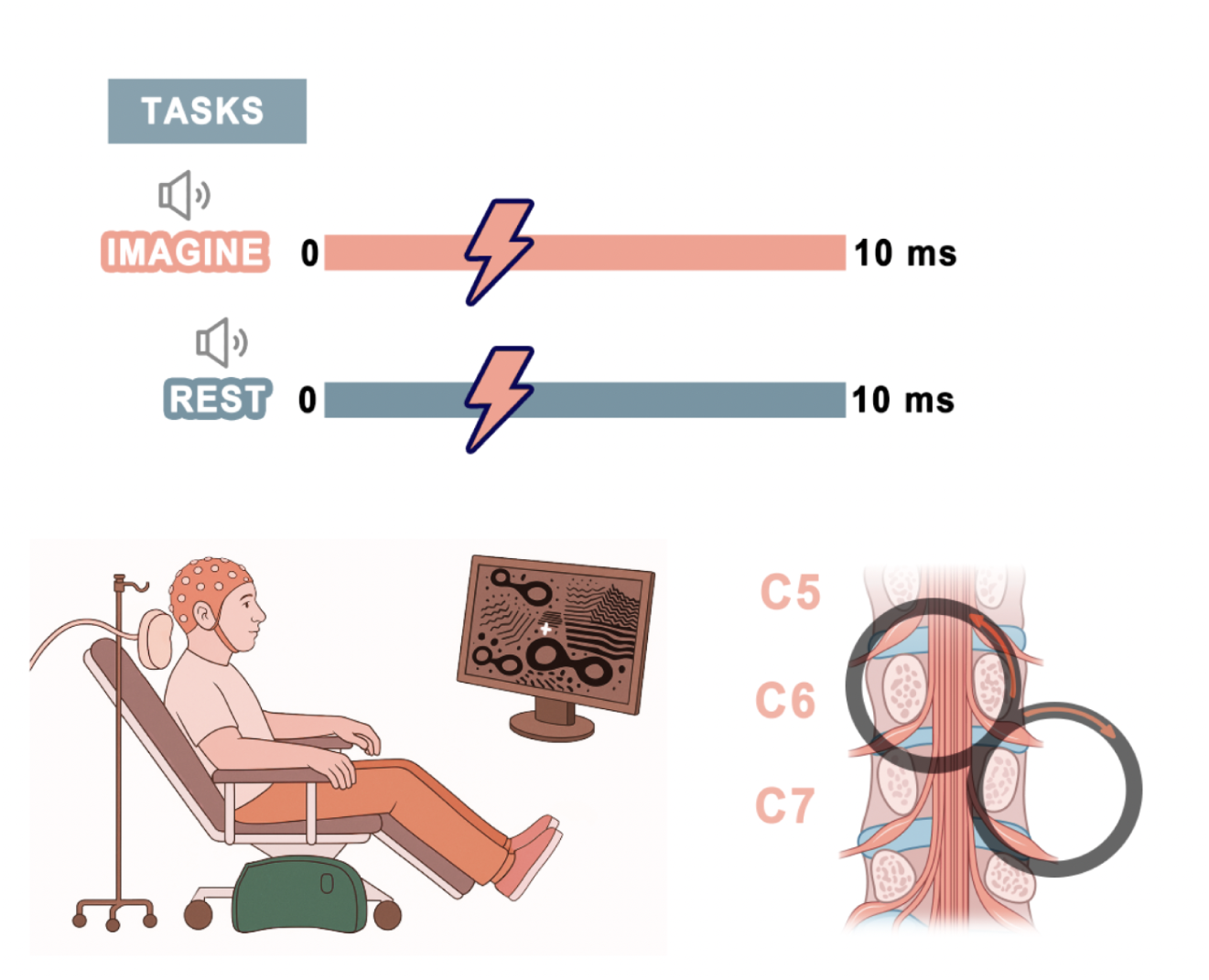
Experimental design. (A) Behavioral tasks. (B) Schematics of the spinal-cord stimulation with a figure-8 coil. Participants were seated comfortably in a TMS chair with their eyes open and gaze fixed on a cross displayed on a screen. An auditory cue signaled them to begin MI or to remain at rest. Participants performed kinesthetic imagery of the right-hand palm closing and opening. Each trial lasted 10 s. The TMS pulse was delivered randomly 4 to 6 s after the start of the auditory signaling to start the imagery. A total of 40 pulses were applied for each condition.

### EMG Recording

Surface electromyographic (EMG) activity was recorded in four muscles: the right flexor digitorum superficialis (R_FDS), abductor digiti minimi (R_ADM), extensor digitorum communis (R_EDC), and the left flexor digitorum superficialis (L_FDS). EMG data were collected during all experimental conditions for further analysis. EMG was sampled at 5 kHz using the Nexstim EMG system.

### Experiment 2

In the second experiment, 20 volunteers (8 females, 12 males), aged 20 to 35 years (mean = 25.3, all right-handed), were recruited under the same inclusion and exclusion criteria as in Experiment 1. All participants gave informed consent prior to participation.

### Experimental Design and Setup

Experiment 2 required participants to perform the same tasks as in experiment 1. Here, single pulse stimulation was applied over the right motor cortex (M1) and the right side of the spinal cord (R_TSMS). The experimental design was identical to experiment 1. The number of magnetic pulses/trials was 70 per condition. Coil position and orientation were controlled using the Nexstim navigation system (NBS) for both cortical and spinal-cord stimulation.

In addition to EMG recordings, electroencephalography (EEG) signals were recorded to investigate cortical responses to TSMS and TMS and their modulation by MI. TMS pulses were delivered using a Nexstim figure-eight coil, oriented at a 45-degree angle with respect to the sagittal plane. Neuronavigation was guided by the individual MRIs. The precentral area in the right-hand knob was targeted. A hot spot for FDS was first found, followed by RMT determination. For the experimental conditions, 120% of RMT intensity was used.

The right-side TSMS was organized the same way as in experiment 1. An adapted system was set up to ensure that the coil position remained stable over the spinal cord during stimulation.

### EEG recording

EEG was recorded using a TMS-compatible NeurOne DC amplifier (Bittium Plc, Kuopio, Finland) with 62 passive Ag/AgCl flat electrodes positioned according to the standard 10-05 International System. The ground electrode was positioned on the forehead, and the reference electrode was placed at the FCz position. Electrode contact impedance was maintained below 5 kΩ. EEG recording was synced with TMS stimulus delivery. EEG was sampled at 20 kHz. Participants wore earplugs during the procedure to minimize auditory artifacts from TMS clicks.

### EMG data processing

EMG signals were epoched from -100 ms to +100 ms relative to the stimulation pulse, with the pre-stimulus period (-100 to 0 ms) serving as a baseline for correction. Pre-stimulus EMG activity was assessed by analyzing the first 100 ms of each epoch. Epochs with signal power exceeding the mean + 3 standard deviations for the pre-stimulus period were excluded (on average, 5% of epochs were excluded per participant per condition).

The peak-to-peak amplitude of motor evoked potentials (MEPs) was calculated for each individual epoch as the difference between the maximum and minimum values within a 40 ms analysis window starting 10 ms after the TMS pulse for M1 stimulation and 5 ms after the TMS pulse for spinal stimulation. Mean MEP amplitudes were then calculated for each participant and condition. MEP latency was determined as the time of the strongest negative deflection in the EMG signal following the magnetic pulse for each individual epoch. Mean MEP latency was subsequently calculated for each participant and condition.

For visualization purposes and to account for differences in signal amplitude between muscles, MEPs were normalized to the peak amplitude obtained during the corresponding rest condition. This enabled the presentation of TSMS and TMS-evoked MEPs in combined figures.

### EEG preprocessing pipeline

In the EEG analysis, EEG data were first downsampled from 20 to 10 kHz and epoched from -1000 ms to 1000 ms relative to the TMS pulse onset. The epochs with artifacts (approximately 2% of the epochs) were removed. Next, the signals within the interval -2 ms to 15 ms relative to TMS onset were replaced by interpolated values calculated using segment-wise linear interpolation (from the scipy.interpolate 1.10.1 package).

Independent Component Analysis (ICA) was performed in two stages. ICA-1 targeted artifacts directly related to TMS pulses, including residual stimulation noise and electrical noise decay around stimulation timing; components to exclude were identified visually and then excluded. On average, 7 components were deleted for each recording at this stage. A notch filter was applied to the cleaned data before running the second ICA (ICA-2).

The ICA-2 aimed at detecting and removing the components representing neurophysiological artifacts, including eye movements, blinks, muscle activity, and cardiac artifacts. ICA-2 components to remove were identified visually and then excluded; the weight of excluded components was applied to the data generated after ICA-1.

Next, a notch filter at 50 Hz was applied to suppress power line noise, followed by band-pass filtering (0.1–45 Hz). Noisy channels were marked and removed, then interpolated, followed by a re-referencing to average reference (common average reference), and corrected to the baseline measured for the interval -0.3 to -0.1s relative to the trial start. ICA and filtering were implemented using the MNE Python library (3.11.3). Figure 2 shows the graphic representation of the signal preprocessing pipeline.

**Figure 2.**
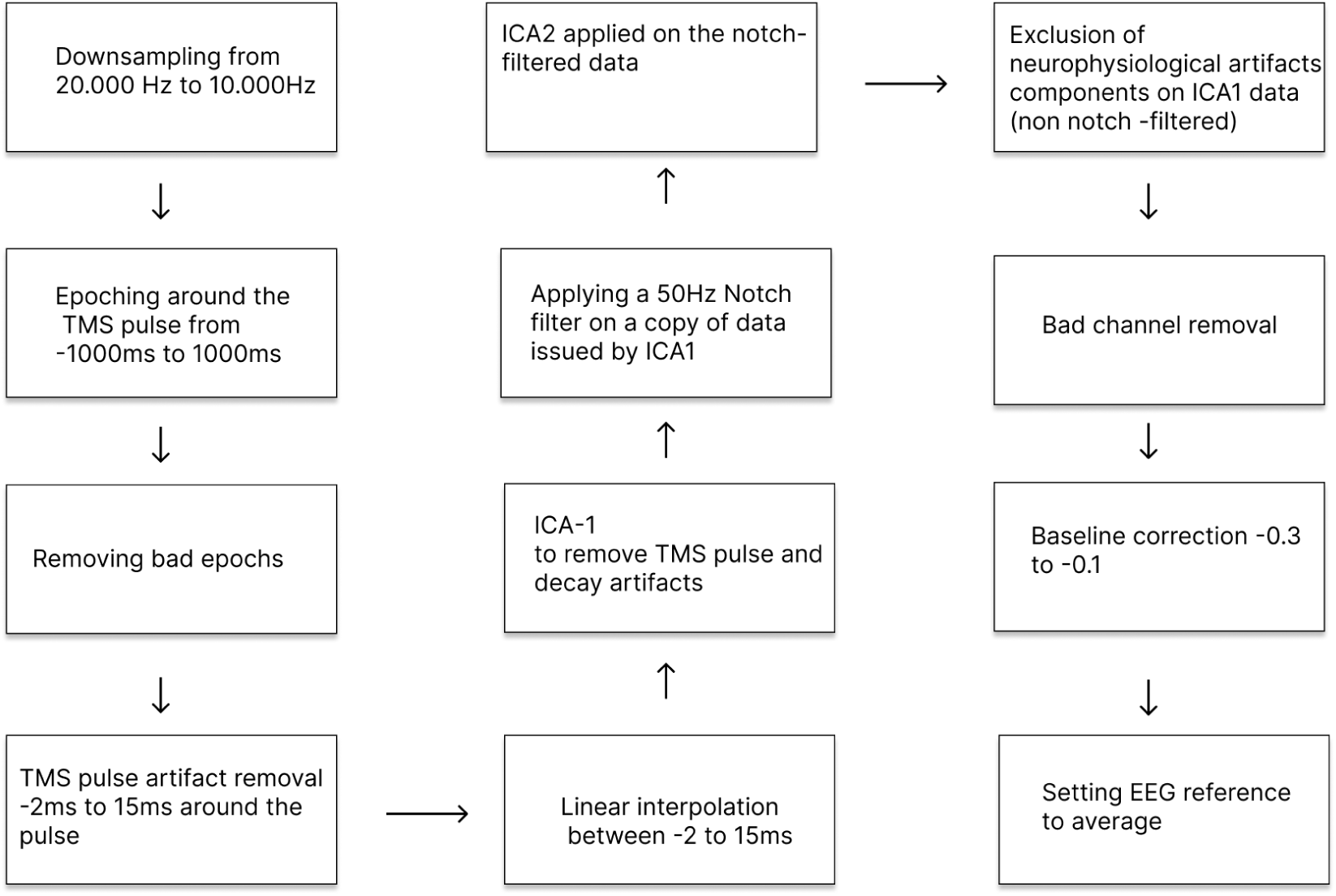
EEG preprocessing steps of, including removal of TMS-related and physiological artifacts and subsequent analysis of TMS-evoked potentials.

### TMS and TSMS evoked potentials

In total, we had N=17 for R_TSMS MI, N=19 for TSMS_Rest, N=19 for M1_MI, and N=18 for MI_Rest files to analyze.

To assess the spatiotemporal dynamics of stimulation-evoked activity, epochs were averaged for each participant across both motor imagery and rest conditions. Grand averages were computed across participants, and topographical maps were generated for different components from 15 ms to 300 ms post-TMS pulse. Time points for the topographical maps were selected based on visual inspection of TEP and TSEP peaks and with reference to the established temporal structure of TMS-evoked potentials reported in the literature (30 ms, 60 ms, 100 ms, 180 ms, 200 ms, 280 ms, and 300 ms).

### Statistics

To identify significant differences in TEPs and TSEPs between MI and rest conditions, a spatiotemporal cluster-based permutation test (Scipy version 1.10.1) was applied to the evoked cortical signals within the 15-350 ms interval (the interpolated time range was excluded from statistical analysis). Statistical comparisons were conducted using a two-sided cluster-based permutation approach with 2000 permutations, using a threshold of *t* = 2.5. Clusters with *p*-values ≤ 0.05 were considered statistically significant. For visualization, mean F-statistic maps were plotted alongside averaged condition-wise waveforms over significant channels.

For MEP statistical analysis, a three-way ANOVA was performed to determine the effects of CONDITION (MI vs. rest), MUSCLE, and AREA (M1 vs. Spine). Post-hoc comparisons were performed using paired two-sided t-tests. Three-way ANOVA with the same set of factors was conducted to analyze MEPs latency changes.

## Results

### Analysis of motor evoked potentials

EMG analysis demonstrated that TSMS selectively activated side-specific motoneuron pools, eliciting unilateral hand contractions. Figure 3A shows the results of experiment 1, where R_TSMS and L_TSMS stimulation evoked muscle activation in the ipsilateral hand. MEP amplitudes exhibited similar latencies and morphology across stimulation sites for all recorded muscles. Moreover, right-hand movement imagery increased right-hand MEP amplitudes in R_TSMS, while MEPs evoked by L_TSMS remained unchanged.

**Figure 3:**
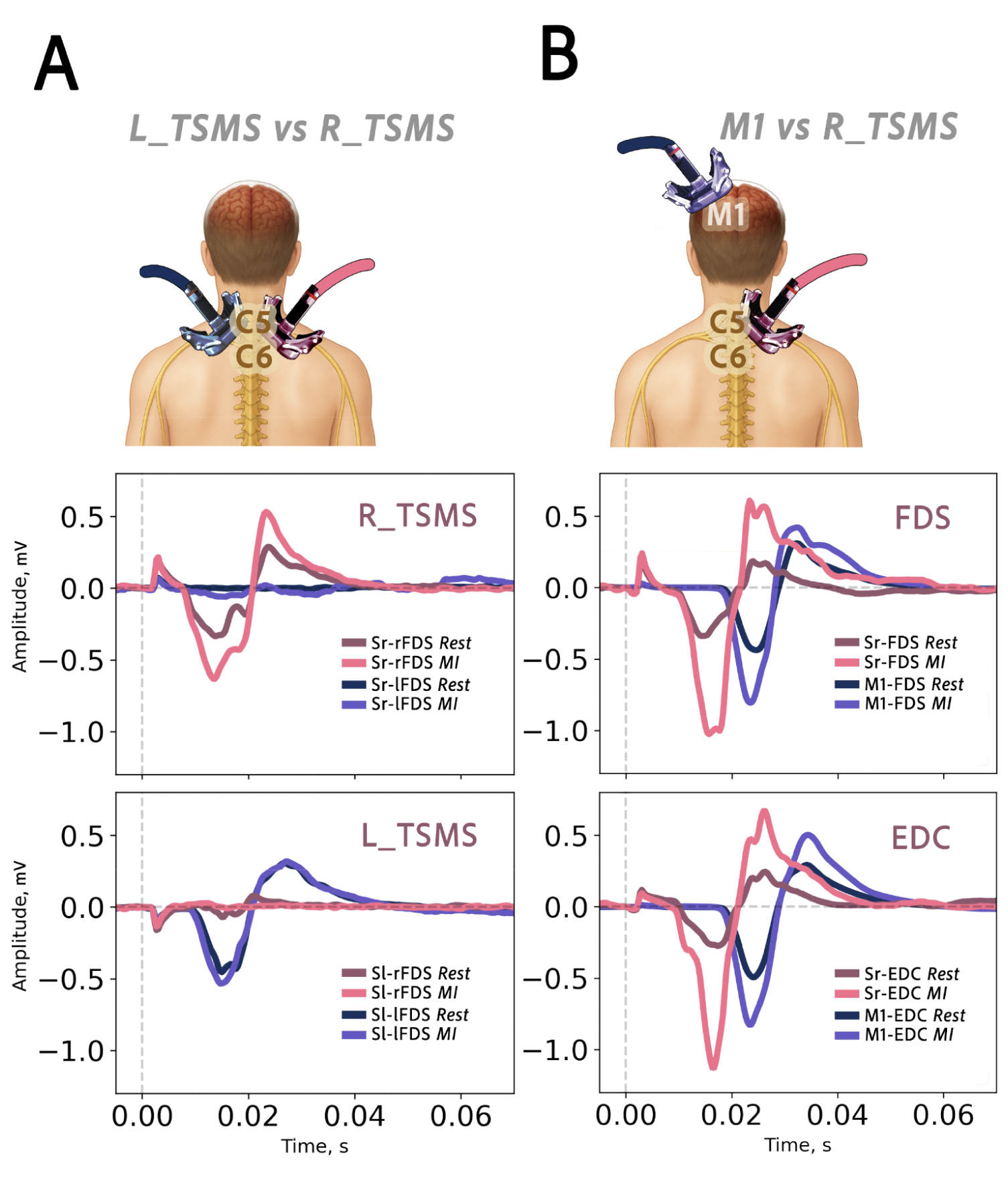
Motor-evoked potentials elicited by spinal (TSMS) and cortical stimulation (TMS). **(A) Experiment 1, (B)** Experiment 2. **R_TSMS:** Right side stimulation of the spinal cord; **L_TSMS:** Left side stimulation of the spinal cord; **M1:** Stimulation over the motor cortex. **FDS:** *flexor digitorum superficialis*, **EDC** : *extensor digitorum communis*. **rFDS**—Right arm FDS, **lFDS**—Left arm FDS.

**Figure 4.**
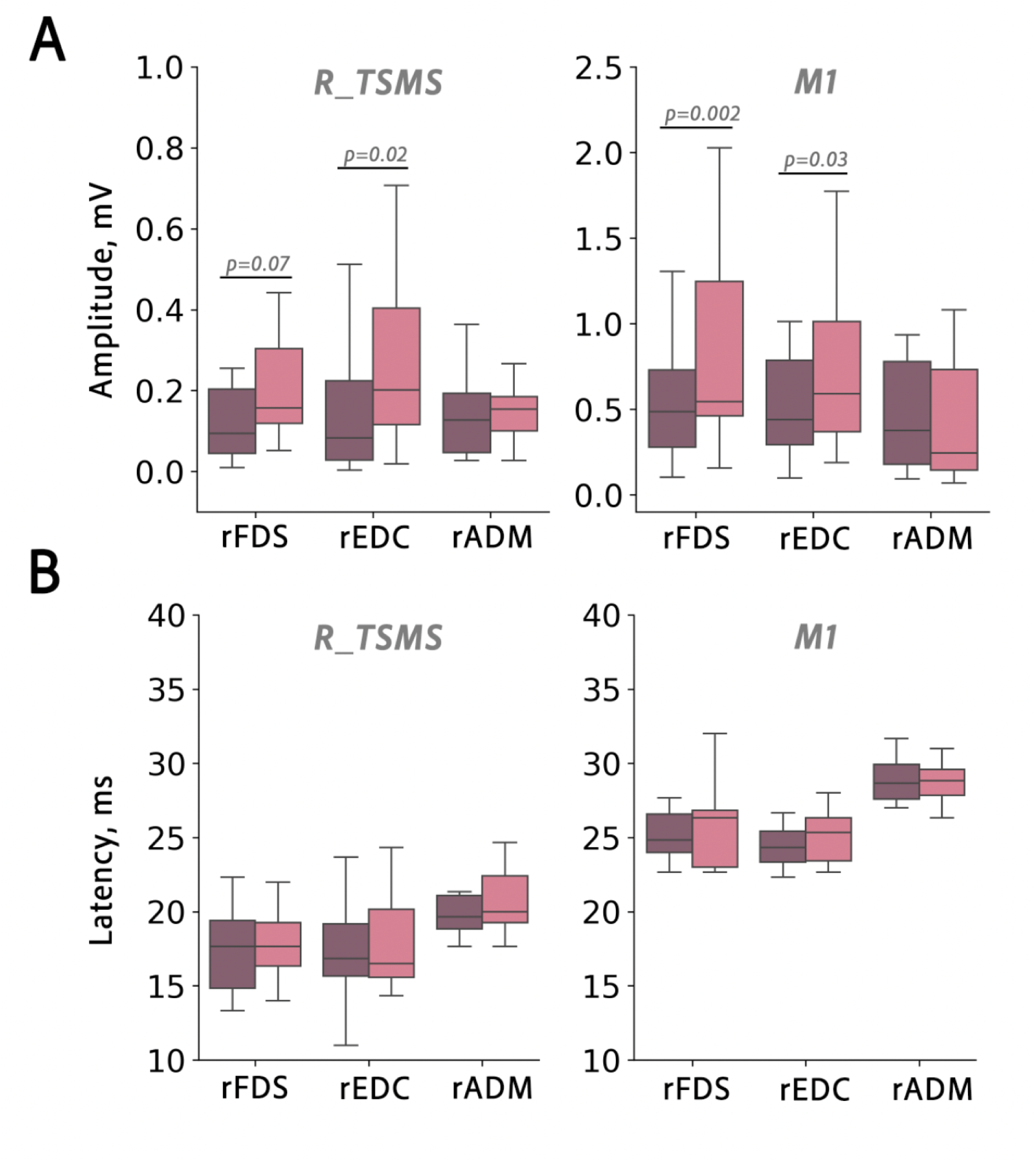
(A) Peak-to-peak amplitudes of the motor-evoked potentials (MEPs) during resting and MI. Data are shown for different muscles and stimulation sites (R_TSMS vs. M1)**. (B) MEP latencies** across conditions, stimulation sites, and muscles.

In the second experiment, we compared motor responses between TSMS and M1 stimulation. Figure 3B demonstrates that both stimulation conditions exhibited increased MEP peak-to-peak amplitudes during MI. A difference in latency was found, where MEPs evoked by TSMS occurred earlier than MEPs evoked by M1 stimulation.

A three-way ANOVA revealed significant effects of condition (MI vs. rest: F(1, 206) = 7.07, p = 0.008, η² = 0.033) and stimulation area (M1 vs. R_TSMS: F(1, 206) = 69.65, p < 0.001, η² = 0.253), indicating larger MEP amplitudes during motor imagery compared to rest and with M1 stimulation compared to TSMS. No significant main effect of muscle (p= 0.203) or interactions (all p > 0.05) were observed.

On the other hand, MEPs latencies showed a significant effect of stimulation site (TSMS vs M1; F = 24.74, p <0.000001, η²= 0.107 ) and significant effect of muscle (F(2, 206) = 11.08, p < 0.001, η² = 0.097), suggesting varying latencies across stimulation sites and muscles [naturally due to physiological anatomy]. The condition ( MI vs Rest ) had no significant effects on MEPs latency (p = 0.231)

### EEG analysis

M1 stimulation-evoked TEP waveforms matched classical TEPs reported in the literature (Beck et al., 2024), exhibiting P30 and P60 peaks followed by N100 and P180 components. Early peaks demonstrated left-sided scalp localization, while later (180-300 ms after stimulus onset) components showed central-parietal distribution (see Fig.5).

**Figure 5.**
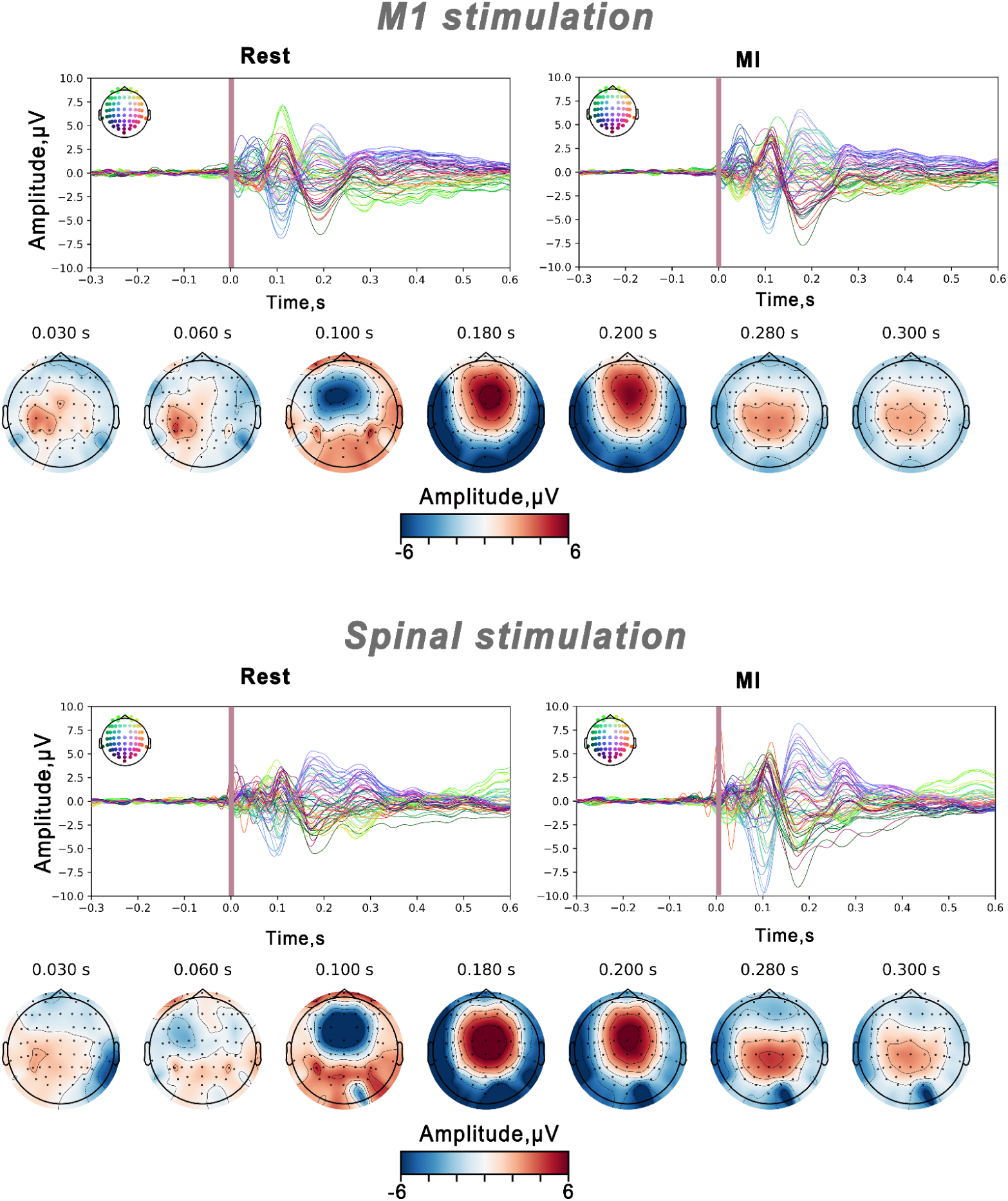
TMS-evoked potentials (TEPs) and TSMS-evoked potentials (TSEPs) across all channels during the rest and right-hand MI conditions. TMS artifact interval is indicated by purple shading. Scalp distribution of specific peaks is shown as topographic maps during MI condition.

The TSMS-evoked cortical potentials exhibited early P30 and, in contrast to M1, a negative deflection at 60 ms N60, with scalp distribution contralateral to the stimulated spinal side. N100 and later components were localized to central-parietal channels.

Cluster-based permutation testing revealed multiple spatio-temporal clusters showing MI effects for both cortical responses. For TEPs, significant clusters included: an early cluster over the stimulated (left) motor cortex at 15–20 ms post-stimulus (N15), two clusters (ipsilateral and contralateral to the stimulation side) corresponding to the P30 component, and a late cluster at ∼100 ms post-stimulus corresponding to the N100 peak. All clusters indicated MI-induced increases in TEPs component amplitudes compared to rest.

For TSEPs, MI also increased component amplitudes. Permutation testing identified an early cluster contralateral to the stimulated spinal side corresponding to the P30 component [23-37ms] around the sensorimotor area, and later significant clusters [70-144ms] and [145-280ms] over centro-parietal and frontal area.

## Discussion

In this study, we used TMS applied to M1 and TSMS to probe modulatory effects of MI upon the corticospinal and spinal excitability within the same experiment. For both TMS and TSMS, we used a figure-of-eight coil. During TSMS it was positioned over the right sixth cervical nerve root (C6, between C6-C7 vertebrae). TSMS delivered in this fashion evoked both ipsilateral muscle responses (i.e., MEPs) and cortical responses associated with the antidromic corticospinal and orthodromic spinocerebral pathways. Right-hand MI increased TSMS-evoked MEPs in a muscle-specific manner. Cortical responses to TSMS were also modulated by MI, as reflected by altered TEPs and TsEPs.

### TSMS evoked MEPs increase during MI

With TSMS, we reliably elicited MEPs using a figure-of-eight coil positioned at the right side of spine between the C6–C7 vertebrae and oriented at ∼45°. Previous studies demonstrated that coil position during TSMS affected the laterality and focality of muscle responses [23]. We confirmed this spatial specificity by showing that by positioning the coil on the right or left side of the spine MEPs could be evoked exclusively in the ipsilateral upper limb (Experiment 1).

Second, MI of the right hand being closed and opened selectively enhanced the R_TSMS-induced MEPs but not the L_TSMS-induced MEPs. This task-specific facilitation can be compared to the literature on the effects of MI on TMS-evoked responses demonstrating that these effects are muscle- or effector-specific [8,10,24–25]. Our results show that such specificity also exists for when responses to TSMS are considered.

In Experiment 2, MI increased muscle responses evoked by both TSMS and M1 TMS, which is consistent with the previously established facilitatory action of MI on corticospinal excitability [8,10,26]. Notably, we observed that TMS evoked stronger MEPs compared to the responses to TSMS. This could be explained by strong corticospinal connectivity for the forearm muscles and possibly weaker spinal reflexes for the same muscles, but also methodological factors could have played a role such as the assurance that TMS stimulated M1 when guided by subject-specific MRI neuronavigation and delivered at 120% RMT, the absence of such assurance for the stimulation of the spinal cord with the TSMS conducted at 110% RMT without MRI-anatomical guidance. Additionally, the relative weakness of the TSMS-evoked responses does not speak against this approach, particularly if TSMS activates different motoneuronal pools compared to TMS. Further investigation of optimal stimulation intensity will be needed for TMS, TSMS, and their combinations.

The analysis of MEP latency showed that TSMS-evoked MEPs had significantly shorter latencies than M1-evoked MEPs (area effect: F = 24.74, p < 1e-6, η² = 0.107). This result is completely expected due to the shorter conduction path for the spinal reflexes evoked by the stimulation of dorsal roots compared to the corticospinal tracts. Differences in muscle-specific motoneuron pool anatomy also contributed to latency variability (muscle effect: F(2, 206) = 11.08, p < 0.001, η² = 0.097). We found that MEP latencies remained unchanged during MI (p = 0.231). While the effects of MI on corticospinal excitability have been reported in many studies, the effects of MI on the spinal circuitry - remain debated. Several studies reported that MI increased spinal reflexes (e.g., H-reflex and F-wave) proportionally to imagined contraction intensity [11, 27–29], which indicates that MI elevated motoneuron excitability to direct inputs [15]. Conversely, other studies of the upper-limb muscles found no MI-related changes in the H-reflexes or F-waves [26, 30]. These mixed findings contrast the overwhelming evidence of MI affecting cortical circuitry that connects to the spinal cord [31].

A plausible explanation for these mixed findings is that classical reflex measures probe restricted spinal circuit elements (monosynaptic Ia-motoneuron pathways for H-reflex; antidromic motoneuron excitability for F-waves) and are strongly influenced by presynaptic inhibitory mechanisms. Consequently, they may not detect modulation of interneuronal, propriospinal, or polysynaptic circuits. TSMS, by contrast, can recruit dorsal roots, ventral roots, and a broader array of intraspinal networks depending on coil geometry and intensity [19,32]. Because we used suprathreshold TSMS with explicit kinesthetic instructions for MI, our paradigm could have engaged these additional spinal elements, enabling detection of MI-related facilitation that monosynaptic reflex protocols might miss [20].

### Cortical responses to spinal and motor cortex stimulation

#### TMS-EEG data pre-processing pipeline

EEG recorded with concurrent TMS or TSMS is highly susceptible to artifacts. Beyond traditional physiological artifacts including muscle activity and eye movements, the magnetic pulse itself induces substantial distortions in the EEG signal, followed by post-pulse decay artifacts. To ensure valid TEPs and TSEPs measurements, we implemented a multi-stage preprocessing pipeline with carefully ordered steps: interpolation around the pulse artifact and two rounds of ICA with component exclusion guided by topography and trial-to-trial variability.

Preprocessing approach and step order markedly influence early TEP characteristics. Several studies confirmed that methodological reporting and preprocessing order vary inconsistently across studies, contributing to variance in TEP metrics [33–39]. Additionally, when identical data were processed through four different pipelines, components before 100 ms showed divergence in amplitude and waveform, while late components demonstrated high inter-pipeline correlation [33]. Recent work by Brancaccio et al. [35] comparing ARTIST, TESA, and SOUND/SSP-SIR using synthetic ground-truth data revealed substantial differences in spatial-temporal precision and inter-trial variability. Therefore, we implemented our pipeline conservatively to preserve early neural activity while minimizing artifact contamination.

### TMS and TSMS evoked potentials

The topography of the TMS-evoked potentials (TEPs) in our study was consistent with the previous reports. The TEPs displayed early components (P30 and P60) localized over the stimulated motor cortex, as well as later components (N100 and P180) with a broader distribution [37–39] during rest and MI conditions.

We observed that MI increased the amplitude of the early TEP components within the 15–32 ms interval after the magnetic pulse (see Figure 6), This aligns with the prior studies reporting that MI enhances cortical excitability [10, 40–42]as the P30 component is considered to reflect early motor cortex activation. The later TEP components displayed bilateral activations, suggesting the involvement of the ipsilateral sensorimotor networks during a unilateral motor task and reflecting a broader cortical engagement that extends beyond the contralateral hemisphere [43–46].

**Figure 6:**
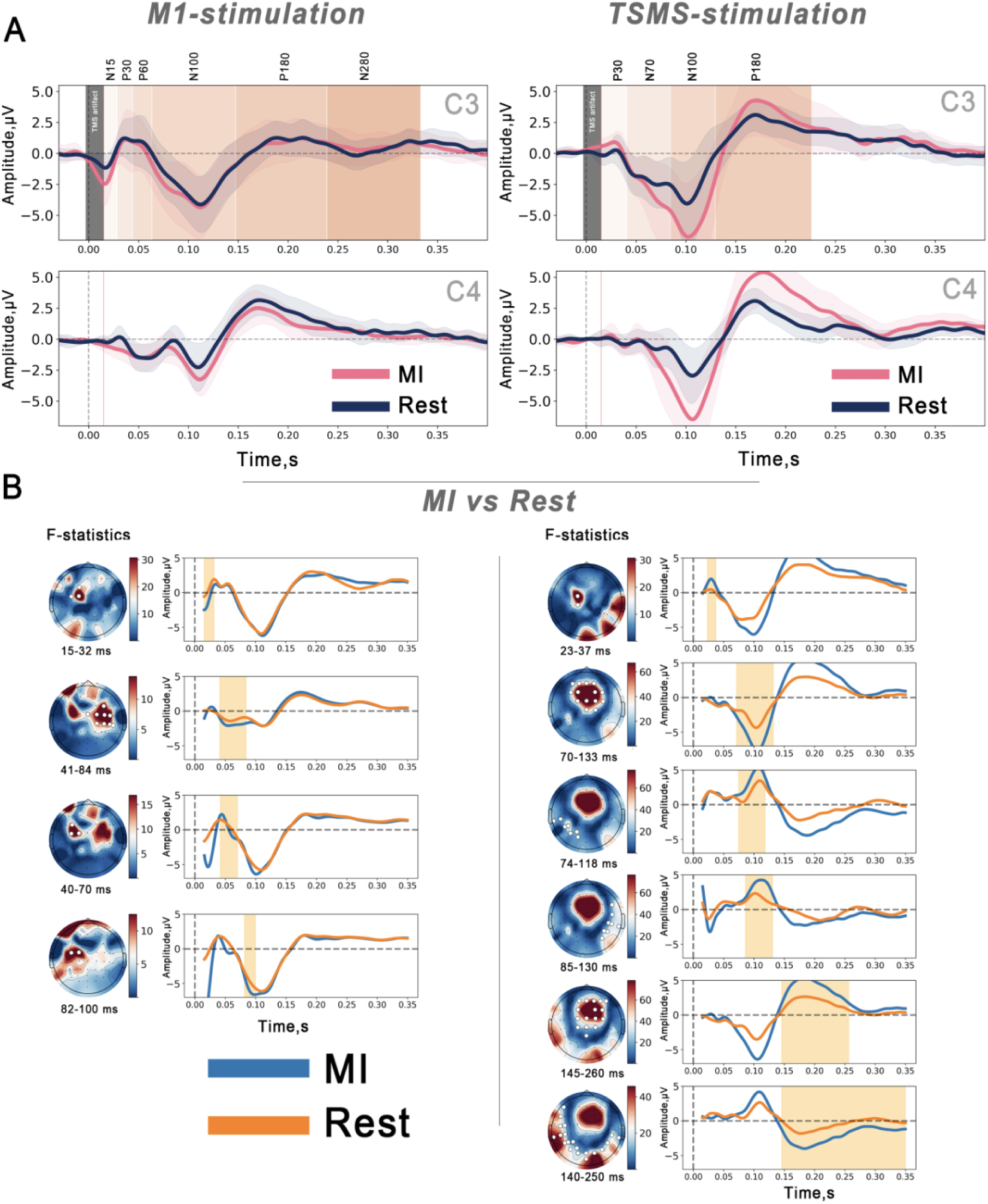
(A) TMS-evoked potentials (TEPs) and TSMS-evoked potentials (TSEPs) over channels C3 and C4 during MI and rest. (B) Cluster-based permutation test showing significant clusters corresponding to TEPs and TSEPs peaks.

Furthermore, we report that TSMS delivered at the cervical level elicits cortical responses (TSEPs) that resemble canonical TEP waveforms evoked by M1 TMS. Early TSEP peaks in the 20–40 ms range may correspond to fast somatosensory afferents and local sensorimotor processing [47–48]. The presence of the P30 component over this area in TSEPs suggests the activation of sensorimotor circuits even when the stimulation pulse originates at the spinal level.

However, a difference emerged in the 60–70 ms time window; where TSEPs showed a negative deflection (N60), whereas M1-evoked TEPs typically display a positive peak at this latency. This polarity inversion could reflect distinct neural processes and pathways engaged by the spinal versus cortical stimulation. Indeed, in M1 TMS studies, the P60 component is thought to reflect afferent input from the stimulated M1 region to the primary somatosensory cortex (S1), likely arising from muscle twitches induced by the TMS pulse and subsequently processed by S1 [39]. Source localization studies have localized the P60 peak to the left superior parietal lobule, associated with S1 processing [43].

However, during TSMS, the magnetic pulse is applied over the cervical spine, which might explain the absence of P60 [50]. Source localization analysis will be needed to determine whether early TSEP components represent purely sensory evoked potentials or also reflect corticospinal network activity and to identify the neural sources activated during different time windows under TSMS condition.

### Motor imagery enhances cortical responses to magnetic spinal stimulation

One key finding in our study is that MI primes the cortex, enhancing its responses to the magnetic spinal stimulation. We found that MI significantly increased TSEPs amplitudes, particularly at the P30 component, indicating that the cognitive priming of the motor cortex enhances early cortical processing of ascending spinal volleys [39, 40,52].

This process might be attributed to the TSMS method generating afferent volleys that ascend to the cortex, activating not only the sensory tracts but also motor neurons and other spinal interneuronal circuits. This aligns with the established mechanisms of other spinal stimulation techniques, known to elicit both sensory potentials and to modulate motor excitability [53–56]. Therefore, our results suggest that MI primes M1 to be more receptive to ascending volleys, resulting in amplified cortical responses over the sensorimotor area after TSMS.

## Novelty

Focal TSMS presents a promising approach for side-specific rehabilitation targeting, potentially enhancing therapeutic outcomes. This study offers several novel insights into the neural mechanisms underlying TSMS. While the effects of motor imagery on MEPs elicited by cortical stimulation are well-documented, this is the first study to investigate the neural correlates of magnetic spinal stimulation during MI using EEG.

Furthermore, TEPs elicited by single TSMS pulses have not been previously characterized. In this study, we meticulously characterized the EEG correlates of TSMS and identified distinct TEP components, revealing similarities and differences in waveform components to those observed in cortical stimulation after rigorous preprocessing.

Finally, by directly comparing TSMS to M1 stimulation under MI conditions, we demonstrate that TSMS responses exhibit different latencies in motor pathways (as measured by EMG and possibly distinct spatial cortical activations).

This study reveals a reciprocal connection between the brain and spinal cord during MI. Indeed MI affects spinal responsiveness to external stimulation, as shown by increased amplitudes and activation patterns, in addition to modulating corticospinal output.

## Limitations and Future Directions

Combining TSMS with MI could collaboratively enhance neuroplasticity by synchronizing cortical and spinal activations, potentially improving recovery in patients with SCI or after stroke. However, further investigation into the neural mechanisms of TSMS is required. Employing source localization techniques and advanced EEG analyses will be crucial to accurately map TSMS-evoked potentials on the cortex and understand their underlying neural substrates.

While our findings suggest that lateralized TSMS coil positioning can evoke ipsilateral responses, we cannot conclusively assert that this specific coil orientation and positioning achieve precise root-specific stimulation without additional position mapping and electrical field modeling studies.

It is also important to note that TEP waveforms can be influenced by peripheral sensory inputs and artifacts. Therefore, meticulous preprocessing and control conditions are essential to isolate genuine cortical responses. In our study, we applied rigorous EEG preprocessing to minimize such confounds; however, further refinement of TMS-EEG pipelines remains an active area of research [33–39].

Lastly, the optimization of simulation parameters for magnetic stimulation of the spinal cord with TMS, including coil orientation and stimulation intensity, requires further investigations to improve the precision and efficacy of TSMS protocols.

## Conclusion

Spinal cord stimulation techniques have demonstrated promising modulatory effects for patients with SCI and stroke, contributing to a decrease in spasticity, alleviation of pain, and improvements in motor functions [57–62]. TSMS emerges as a promising method not only for probing central nervous system excitability but also for serving rehabilitative purposes. The current study provides insights for future rehabilitative protocols exploring the combination of motor imagery with TSMS, leveraging both the top-down priming of the motor cortex by MI and the bottom-up effects of spinal stimulation to the cortex.

